# ELiAH2: update of the atlas of E3 ligases in human tissues by integrating tissue-specificity and docking properties of ligands for targeted protein degradation

**DOI:** 10.64898/2026.10.10.758184

**Authors:** Sangjae Seo, Seunghun Song, Younghoon Kim, Moon Ki Kim, Hyeonjae Lee, Hyojung Paik

## Abstract

Targeted protein degradation (TPD), including proteolysis-targeting chimeras (PROTACs), requires coordinated selection of E3 ligases, target proteins, and ligands to achieve efficient and tissue-selective degradation. However, current TPD resources provide limited support for integrating tissue-specific transcriptomic evidence with structural compatibility during early candidate prioritization. Here, we present ELiAH2, an expanded database for transcriptome- and structure-guided prioritization of TPD candidates. ELiAH2 integrates GTEx RNA-seq profiles across 24 human tissues to characterize tissue-specific expression patterns of E3 ligases and target genes. Compared with the previous version, ELiAH, current update presented over 200% of expansion for the number of covered tissues and samples. Previously, E3 ligase– target gene associations were inferred using mutual information-based network analysis, enabling users to explore transcriptomic relationships. To further support structural evaluation, ELiAH2 provides large-scale docking datasets comprising 827,385 E3 ligase–target protein pairs and 7,800,521 protein–ligand docking models, including E3 ligase–ligand and target protein–ligand interactions. Docking scores, ranked interaction tables, gene and ligand identifiers, and downloadable PDB structures are incorporated into the web interface. By combining tissue specificity, E3–target association, protein–protein docking, and protein– ligand docking, ELiAH2 provides a comprehensive resource for narrowing the TPD search space and supporting rational degrader design. ELiAH2 is freely available at https://eliahdb.org.

## INTRODUCTION

Targeted protein degradation (TPD) has emerged as a promising therapeutic modality for targeting undruggable proteins. Unlike conventional inhibitors, TPD technologies induce selective degradation of target proteins by recruiting endogenous E3 ubiquitin ligases to the ubiquitin–proteasome system (1,2). TPD offers pharmacological advantages over conventional drugs by enabling cyclical degradation events, thereby achieving prolonged target suppression, improved efficacy and targeting of undruggable proteins (3). Consequently, TPD has rapidly evolved from a proof-of-concept technology into a major drug discovery platform, with multiple degrader molecules currently undergoing clinical evaluation.

The success of TPD critically depends on the selection of an appropriate E3 ubiquitin ligase. Although more than 600 human E3 ligases have been identified, only a limited number, such as CRBN, VHL, MDM2, and cIAP1, have been widely exploited for degrader development (4–6). Increasing evidence suggests that the expression abundance, tissue specificity, and molecular compatibility of E3 ligases substantially influence degradation efficiency, therapeutic selectivity, and potential off-target effects (7,8). Since the recruited E3 ligase and target protein must coexist in the same cellular environment, tissue-specific transcriptomic profiles are essential for rational degrader design. Furthermore, molecular level interaction data based on tissue-specific transcriptomic profiles could be an important basis for rational degrader design. Nevertheless, the incorporation of transcriptomic evidence into E3 ligase and target protein prioritization remains limited.

Several publicly available resources have facilitated TPD research by collecting experimentally validated degraders, E3 ligase–substrate interactions, and chemical information. For example, PROTAC-DB provides comprehensive annotations of reported degrader molecules and their pharmacological properties (9), whereas UbiNet curates experimentally supported E3 ligase– substrate interactions (10). More recently, E3Atlas systematically characterized the expression landscape of human E3 ligases across normal tissues (11). Despite these valuable resources, most lack an integrated framework that combines tissue-specific transcriptomic data, E3 ligase–target regulatory associations, and structure-based interaction analysis for tissue-aware E3 ligase prioritization.

To overcome these limitations, we previously developed ELiAH (E3 Ligase Atlas in Human Tissues) (12), a transcriptome-based database that systematically characterized tissue-specific expression patterns of E3 ligases and their associated genes using GTEx RNA sequencing data. ELiAH enabled users to explore tissue-specific E3 ligases and infer transcriptional associations that may support rational target prioritization for TPD development. However, the first version was limited by its relatively small number of analyzed tissues and lacked comprehensive structural information that could support molecular-level interpretation of E3 ligase–target compatibility.

Here, we present ELiAH2, a substantially expanded database for transcriptome-guided targeted protein degradation. Compared with the previous release, ELiAH2 incorporates a considerably larger collection of GTEx transcriptomes covering 24 human tissues, including multiple brain regions and liver tissue, thereby providing a more comprehensive representation of tissue-specific transcriptional landscapes. Additionally, ELiAH2 incorporates large-scale docking calculations to evaluate E3 ligase–target, E3–ligand, and target–ligand interactions. By combining expression profiles, regulatory associations, and structural compatibility within a single resource, ELiAH2 provides a comprehensive framework for prioritizing tissue-appropriate E3 ligases and target proteins, facilitating the rational design of next-generation targeted protein degraders.

## MATERIALS AND METHODS

### Data collection and RNA sequencing analysis

RNA sequencing (RNA-seq) data were obtained from the Genotype-Tissue Expression (GTEx) (13) Project (version 2.0) through dbGaP under authorized access. Rather than using preprocessed expression matrices, raw FASTQ files were analyzed to maintain a consistent computational workflow and ensure standardized quality control across all samples. In total, 9,247 RNA-seq datasets representing 24 normal human tissues were initially collected.

To minimize technical variation, sequencing quality was assessed using HISAT2 (14) alignment statistics. Samples exhibiting an overall alignment rate below 80% were excluded from downstream analyses. Transcript abundance was quantified using Salmon (15) with transcript-level quasi-mapping, and gene-level expression values were summarized using the GENCODE reference annotation. Sample-wide normalization was subsequently performed using the DESeq2 (16) median-of-ratios method to reduce sequencing depth bias and improve cross-sample comparability.

### Identification of tissue-specific genes and E3 ligases

A comprehensive list of human E3 ubiquitin ligases was compiled from publicly available annotations and manually curated reference datasets. Following normalization, tissue-specific expression was evaluated for every protein-coding gene, including all curated E3 ligases. Statistical significance of tissue-specific expression was assessed using the one-tailed Mann– Whitney U test by comparing expression levels in one tissue against all remaining tissues. Multiple hypothesis testing was corrected using the Benjamini–Hochberg procedure, and genes with an adjusted false discovery rate (FDR) below 0.05 were regarded as tissue-specific.

### Inference of E3 ligase–target gene associations

Potential regulatory associations between E3 ligases and protein-coding genes were inferred using ARACNe-AP (Algorithm for the Reconstruction of Accurate Cellular Networks through Adaptive Partitioning) (17). Mutual information (MI) was calculated between each E3 ligase and every expressed gene across all analyzed samples.

To improve network robustness, ARACNe-AP performed multiple bootstrap iterations followed by consensus network reconstruction. Statistical significance of each inferred interaction was estimated from the frequency of edge detection across bootstrap networks using a Poisson distribution model. P-values were corrected for multiple testing using the Bonferroni method. Significant E3 ligase–target gene associations were incorporated into the ELiAH2 database together with corresponding mutual information scores.

### Integrated structural docking analysis of protein–protein and protein–ligand interactions

To evaluate structural compatibility among E3 ligases, target proteins, and ligands, ELiAH2 performed integrated docking analyses for both protein–protein and protein–ligand interactions. Three-dimensional protein structures were retrieved from the AlphaFold Protein Structure Database by querying gene symbols and mapping them to the corresponding UniProt entries. When available, AlphaFold-predicted PDB structures were downloaded and used as input structures for subsequent docking analyses.

For E3 ligase–target protein interaction analysis, global protein–protein docking simulations were performed using RosettaDock. In total, 827,385 E3 ligase–target protein pairs were subjected to protein–protein docking analysis. For each pair, multiple decoy structures were generated, and the model with the lowest Rosetta interface score (I_sc) was selected as the representative docking model. The corresponding interface score and PDB structure were incorporated into the ELiAH2 database. Lower interface scores indicate more favorable protein–protein interactions and were used as complementary structural evidence for prioritizing candidate E3 ligase–target pairs.

For protein–ligand docking analysis, ligand binding affinity was independently calculated for E3 ligase–ligand pairs and target protein–ligand pairs. The same AlphaFold-derived protein structures were used as receptor structures, and ligand structures were retrieved from the PROTAC-DB 3.0. Ligand binding poses were first predicted using DiffDock, and the resulting protein–ligand complex structures were subsequently evaluated using RosettaDock. In total, 282,540 E3 ligase–ligand pairs and 7,517,981 target protein–ligand pairs were analyzed. For each protein–ligand pair, multiple docking models were generated, and the structure with the lowest I_sc was selected as the representative docking model. The final docking scores and corresponding PDB structures were stored in the ELiAH2 database and made available through the web interface.

## RESULTS

### Database overview and expanded data contents

ELiAH2 was developed to provide a comprehensive transcriptome-guided resource for prioritizing E3 ligases in targeted protein degradation (TPD). Compared with the previous release, the updated database substantially expands both transcriptomic coverage and structural annotations by integrating RNA sequencing profiles from 13 human tissues together with large-scale docking analyses (Figure 1).

**Figure1.**
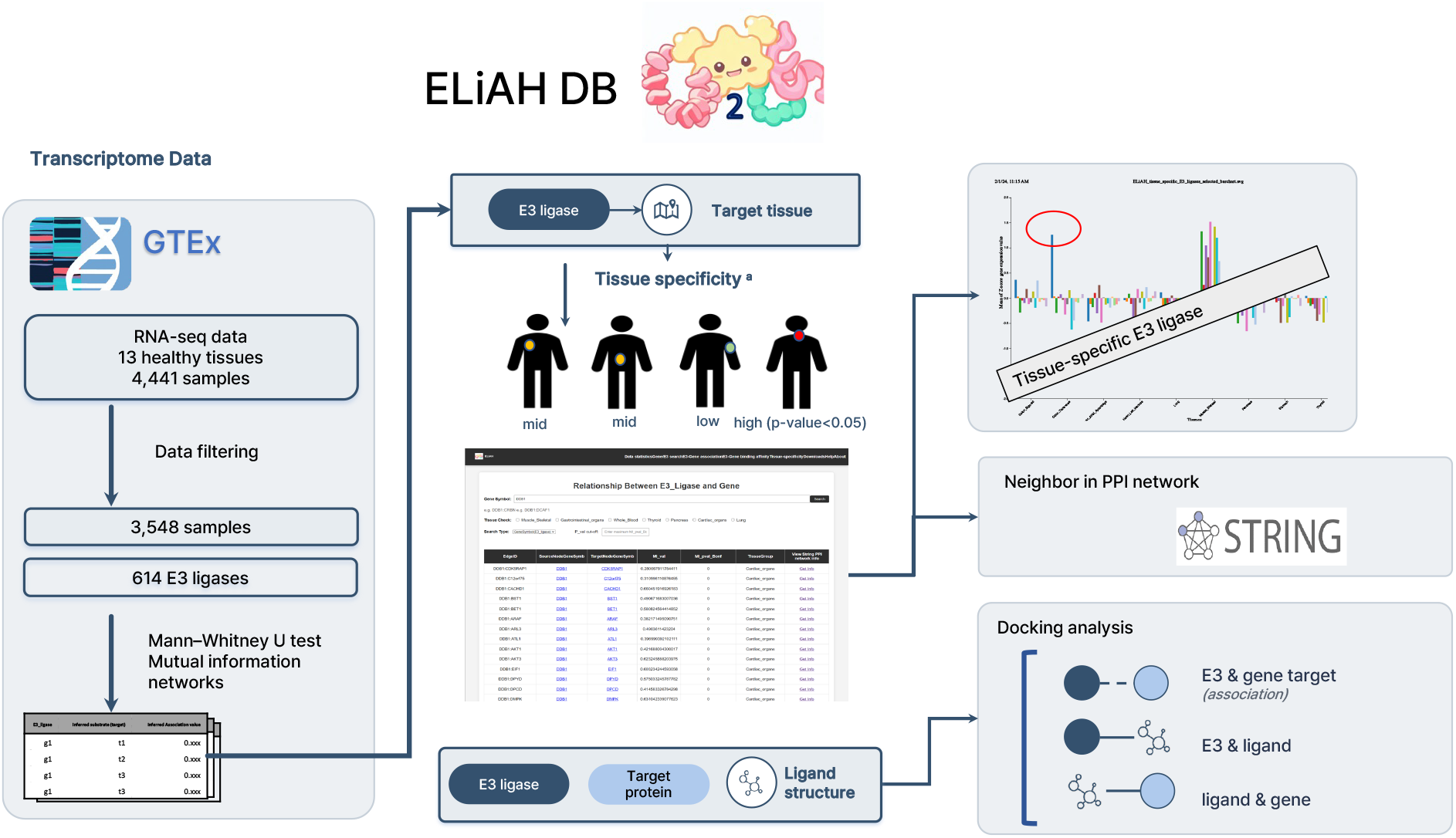
Overview of ELiAH2

Starting from 9,247 GTEx RNA-seq samples, stringent quality-control procedures yielded 3,548 high-quality transcriptomes for downstream analysis (Table 1). These samples encompass multiple physiologically distinct organs, including gastrointestinal tissues, skeletal muscle, thyroid, lung, pancreas, liver, blood, cardiac tissues, and five anatomically separated brain regions. The expanded tissue coverage enables users to investigate tissue-dependent transcriptional characteristics with substantially improved anatomical resolution compared with the previous version.

**Table 1.** Statistics of RNA-seq data used in ELiAH2.

| Data | Frequency |
| --- | --- |
| Total number of samples in GTEx (ver. 2.0) | 9,247 |
| Number of analyzed samples (24 tissues) | 4,441 |
| Number of utilized samples (mean of alignment <sup>a</sup> 93.4% ± 3.17% ) | 3,548 |
| Gastrointestinal organs (Stomach, Colon - Transverse, Sigmoid) | 544 |
| Cardiac organs (Left ventricle, Atrial appendage) | 411 |
| Muscle – Skeletal | 366 |
| Whole Blood | 291 |
| Thyroid | 272 |
| Lung | 225 |
| Pancreas | 183 |
| Liver | 134 |
| Brain - Basal ganglia (Brain - Caudate, Putamen, Nucleus accumbens) | 294 |
| Brain - Cortical areas<br>(Brain - Cortex, Frontal Cortex (BA9), Anterior cingulate cortex (BA24)) | 287 |
| Brain - Cerebellum (Brain - Cerebellar Hemisphere, Cerebellum) | 216 |
| Brain - Diencephalon & Limbic System<br>(Brain - Amygdala, Hippocampus, Hypothalamus) | 206 |
| Brain - Central & Medulla<br>(Brain - Substantia nigra, Spinal cord (cervical c-1)) | 119 |

<sup>a</sup> The mean of alignments was measured using HISAT2.

Using the updated transcriptomic dataset, ELiAH2 systematically quantified the expression profiles of 614 curated human E3 ubiquitin ligases together with all expressed protein-coding genes. Tissue-specific expression patterns were evaluated using non-parametric statistical analysis, providing a comprehensive atlas of E3 ligase abundance across normal human tissues.

### Tissue-specific landscape of E3 ligases and target genes

The expanded transcriptomic dataset enabled a more comprehensive evaluation of tissue-specific transcriptional programs than was possible in the previous release. Principal component analysis demonstrated clear separation of tissue groups based on highly variable genes, confirming that the selected transcriptomes preserve biologically meaningful tissue identities.

For each tissue, tissue-specific genes were identified using the Mann–Whitney U test with false discovery rate correction. The resulting expression atlas allows users to examine both E3 ligases and potential target genes under identical statistical criteria. Interactive visualization modules provide normalized expression profiles across all analyzed tissues, allowing direct comparison of candidate E3 ligases before degrader design.

The inclusion of liver tissue and anatomically resolved brain regions substantially increases the biological utility of ELiAH2. Because many therapeutic targets exhibit tissue-restricted expression patterns, this expanded coverage facilitates more accurate prioritization of tissue-compatible E3 ligases while helping anticipate possible degradation in non-target tissues.

### Genome-wide inference of E3 ligase–target gene associations

To characterize transcriptional relationships between E3 ligases and their potential substrates, mutual information networks were reconstructed using ARACNe-AP. Unlike simple correlation analysis, the mutual information framework captures both linear and non-linear transcriptional dependencies, thereby enabling more comprehensive identification of biologically associated gene pairs.

ELiAH2 contains statistically significant E3 ligase–target gene associations inferred from consensus network reconstruction. For each interaction, users can retrieve mutual information scores, statistical significance, tissue-specific expression profiles, and corresponding protein– protein interaction annotations obtained from the STRING database.

This integrated framework enables simultaneous assessment of transcriptional abundance, tissue specificity, regulatory association, and functional interaction, allowing users to prioritize candidate degradation pairs using multiple complementary sources of evidence rather than relying solely on expression levels.

### Large-scale structural interaction landscape for TPD candidate prioritization

To extend ELiAH2 from a transcriptome-based E3 ligase atlas to a structure-informed TPD prioritization platform, we generated large-scale docking datasets for E3 ligase–target protein pairs and protein–ligand pairs. The current ELiAH2 release includes docking results for 827,385 E3 ligase–target protein pairs, 282,540 E3 ligase–ligand pairs, and 7,517,981 target protein–ligand pairs. These datasets were integrated into the database together with docking scores, rank information, gene identifiers, ligand identifiers, and downloadable PDB structures. For E3 ligase–target protein docking, Rosetta interface scores showed a broad distribution (Figure 2(a)), indicating substantial variability in predicted structural compatibility among candidate degradation pairs. Because interface scores scale with interface size and are not directly comparable across structurally diverse pairs, ELiAH2 provides ranked docking results for each target protein rather than a single global cutoff. For example, for target protein HIP1R, the top-ranked E3 ligases by docking score were HIP1R–ASB11 (−33.14), HIP1R–KCNV1 (−31.45), and HIP1R–RNF26 (−27.91), illustrating that structurally favorable E3–target combinations can be identified beyond the most commonly used degrader-recruiting ligases such as CRBN and VHL.

**Figure2.**
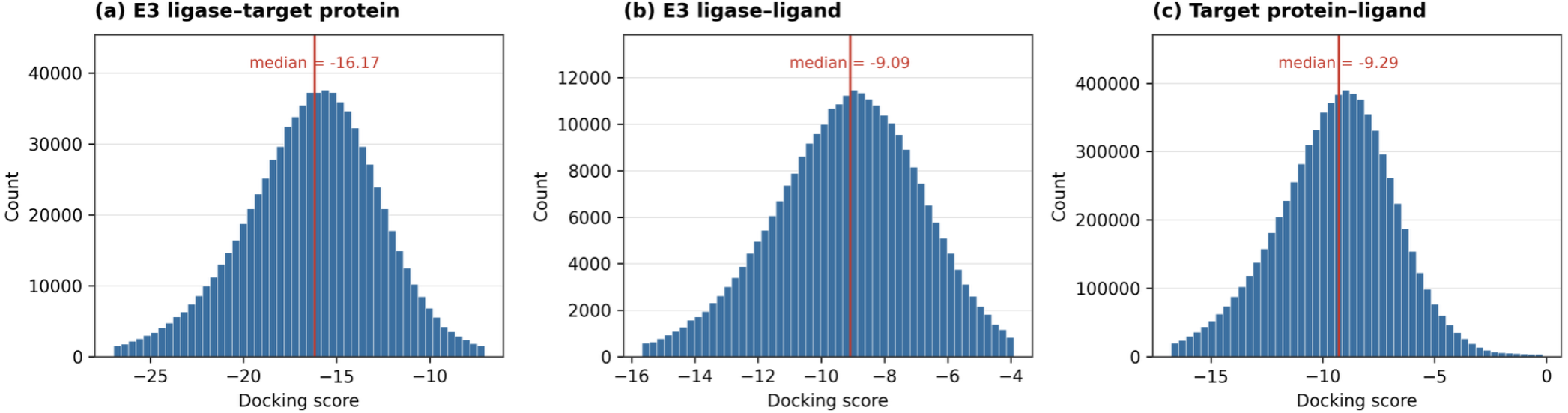
Distribution of representative docking scores for (a) E3 ligase–target protein, (b) E3 ligase–ligand, and (c) target protein–ligand pairs. Red line indicates the median score for each interaction type.

In addition to protein–protein docking, ELiAH2 also provides large-scale protein–ligand docking results for both E3 ligase–ligand and target protein–ligand interactions. A total of 633 unique ligands were docked against E3 ligases and target proteins, yielding 282,540 E3 ligase– ligand pairs and 7,517,981 target protein–ligand pairs (Figure 2(b, c)). As shown by the resulting score distributions, ligands can be ranked according to their relative binding affinities toward a given E3 ligase or target protein. This ranking can be used to propose candidate ligands that can serve as E3 ligase binders or target protein warheads in PROTAC design.

## CONCLUSIONS

ELiAH2 provides an expanded and integrated resource for transcriptome-guided targeted protein degradation (TPD) research by systematically combining tissue-specific gene expression, E3 ligase–target gene associations, and structure-based interaction analyses within a single platform. Compared with the previous release, ELiAH2 substantially increases transcriptomic coverage by incorporating GTEx RNA-seq data from 24 human tissues, enabling more comprehensive characterization of tissue-specific E3 ligases and their potential target proteins. The integration of mutual information-based regulatory networks with large-scale structural docking analyses further allows users to prioritize candidate E3 ligase–target pairs using complementary transcriptomic and structural evidence.

To perform systematic structural evaluation at scale, we employed the Nurion supercomputer and conducted high-throughput, massively parallel docking computations, enabling the completion of over 8.6 million structural docking runs across E3 ligase–target protein and protein–ligand interactions. Together with interactive visualization of tissue-specific expression profiles, regulatory associations, protein–protein interaction networks, and docking results, ELiAH2 provides a practical decision-support platform for selecting appropriate E3 ligases during the early stages of degrader development.

Future updates will further expand the database by incorporating additional transcriptomic datasets, experimentally validated degrader information, proteomics-based evidence, and single-cell transcriptomic profiles to improve cell type-specific prioritization of E3 ligases. Continuous integration of emerging structural prediction methods and experimentally determined ternary complex structures will further enhance the predictive capability of the database.

In summary, ELiAH2 represents a comprehensive and freely accessible database for tissue-aware targeted protein degradation research. By integrating multi-level molecular information into a unified analytical framework, ELiAH2 facilitates rational selection of E3 ligases and target proteins, thereby supporting the development of safer and more effective TPD therapeutics.

## DATA AVAILABILITY

### Accession of web database

ELiAH2 is freely accessible online data available at: https://eliahdb.org/. Users can access this resource without registration or login. The raw RNA-seq data (phs000424.v2 dbGAP) of GTEx consortium can be shared with permission of GTEx consortium.

## SUPPLEMENTARY DATA

Supplementary Data, Supplementary Figure are available at *Database* online.

## AUTHOR CONTRIBUTIONS

Sangjae Seo: Conceive the main idea, Carried out docking property simulation and draft writing

Seunghun Song: Carried out docking property simulation

Younghoon Kim: Review and supporting of draft writing

Moon Ki Kim: Review and supporting of draft writing

Hyeonjae Lee: Review and supporting of draft writing

Hyojung Paik: Conceive the main idea, data acquisition, bioinformatic analysis design, supervise overall project, and funding acquisition.

## ACKNOWLEDGEMENTS

This research was supported by the National Research Council of Science & Technology (NST) grant by the Korea government (MSIT) (No. GTL24033-000). This work also was supported by the National Research Foundation of Korea (NRF) grant funded by the Korea government (MSIT) (No. RS-2023-00257666). This work was supported by the National Supercomputing Center with supercomputing resources including technical support KSC-2025-CHA-0024. This work was supported by the National Research Foundation of Korea (NRF) grants funded by the Korea government (MSIT) (RS-2025-16065483; RS-2025-02315385). HP, HL and YK were also supported by the project of Building and Operating the Korea-Bio Data Station (K-BDS) Platform building and tool development (NRF-2021M3H9A2030520). This work was supported by the Korea Institute of Science and Technology Information (KISTI) (K26L1M2C3).

